# waveome: a toolkit for longitudinal omics analysis using Gaussian processes

**DOI:** 10.64898/2026.03.03.709362

**Authors:** Allen Ross, Jason Lloyd-Price, Ali Rahnavard

## Abstract

Identifying meaningful associations from small-sample longitudinal data is challenging, especially in low signal-to-noise environments where the Gaussian likelihood assumption does not hold. We introduce two methods to algorithmically perform variable selection with sparse, irregularly sampled, longitudinal count data with over-dispersion to characterize nonlinear relationships between omics measurements and covariates of interest using Gaussian processes. The first is an additive non-greedy search-based method, while the second is a penalization approach using Horseshoe priors on kernel hyperparameters. In simulation studies, both methods outperform conventional statistical models in terms of distributional fit and exhibit a trade-off in feature selection. Applying the penalized variant to a real-world Crohn’s disease cohort, we recover well-established biomarkers, such as short-chain fatty acids, secondary bile acids, and specific lipid species, and uncover novel candidates for cross-sectional and temporal disease severity. Both methods are implemented in an open-source Python library, `waveome`, offering a robust set of tools for longitudinal biomarker discovery.

## 1 Introduction

Understanding and characterizing temporal dynamics is essential for studying biological systems. Specifically in biomedical research, the identification of biomarkers that can distinguish disease states is crucial to understanding their etiological mechanisms. This task is particularly challenging for chronic conditions marked by cycles of remission and relapse, due to their inherently temporal patterns of fluctuating disease activity. The importance of this is underscored by the growing global burden of these diseases over the past two decades [1–3]. Consequently, researchers need flexible analytical tools capable of handling repeated measurements from individuals over time. Key characteristics of these data include within- and between-individual variability, potentially nonlinear temporal effects, small sample sizes, and irregular sampling intervals.

These challenges are amplified in omics studies, such as metabolomics, due to low signal-to-noise ratios and non-Gaussian likelihoods [4]. Gaussian processes (GPs) have shown promise in addressing these complexities in prior studies; here, we develop a framework using GPs to model metabolomic intensities and associated metadata, with a particular emphasis on temporal dynamics and variable selection. By integrating cutting-edge GP techniques with variable selection strategies, our approach flexibly filters and ranks candidate biomarkers relevant to a given hypothesis. We have implemented these methods in a Python library, `waveome` (see Figure 1) named from the generally nonlinear relationships in longitudinal omics data, and made it publicly available to the biomedical community.

**Fig. 1:**
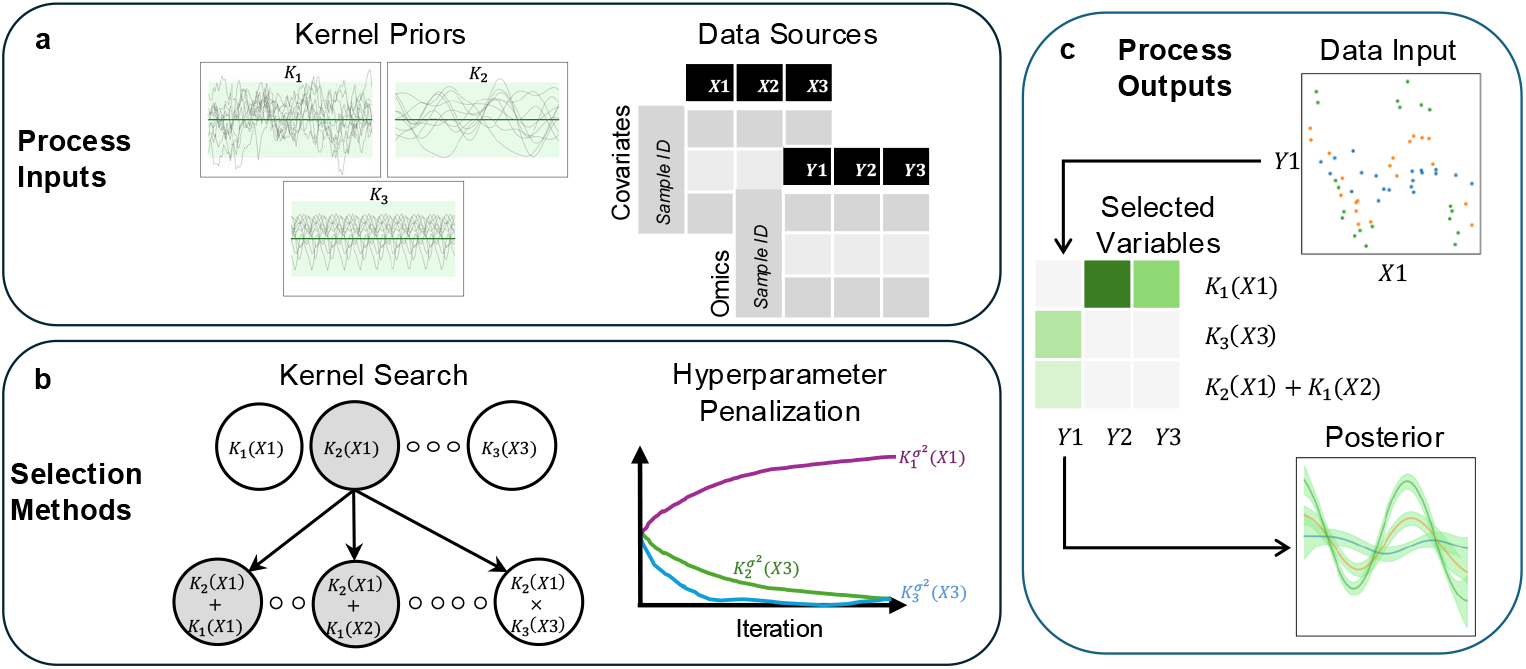
Overview of `waveome` process. **a** Inputs include desired functional forms (kernel similarity functions) to test for each covariate as well as two datasets, both at the sample level: covariates and the omics values. **b** Two options for variable selection methods 1) kernel search through combinations of kernels or 2) penalization of a fully saturated model based on kernel-covariate pairs, where each independent model uses a single omic column as the dependent variable. **c** The output includes a Gaussian process model for each omic feature with selected kernels, covariates, and association strength metrics. Results can be summarized in a heatmap or examined at the level of individual models.

Statistical techniques for longitudinal data analysis are broadly divided into parametric and nonparametric frameworks. Parametric methods typically rely on generalized linear mixed effects models to account for within-unit correlation, accommodate non-Gaussian outcomes, and yield valid inference for fixed and random effects [5, 6]. Recent extensions have integrated variable selection mechanisms, such as penalized likelihood and gradient-based procedures, to enhance model interpretability and performance in high-dimensional settings [7–10].

Non-parametric approaches offer complementary flexibility by avoiding explicit functional assumptions. Local regression and global smoothing techniques, implemented via kernel methods or spline bases, can capture complex temporal patterns but often incur substantial computational cost and complicate uncertainty quantification [11]. GP models occupy an intermediate position: they constitute the Bayesian analogue of kernel regression and generalize spline smoothing through a fully probabilistic formulation [12]. Adaptations of GPs for longitudinal studies include deep kernel learning to capture hierarchical structure [13], additive GP decompositions for interpretable effects [14], and automated kernel discovery algorithms that infer functional structure from data [15–17]. In addition, penalization schemes have been proposed to perform variable selection within the GP framework [18, 19], and specialized covariance functions have been developed to address the themes of repeated measurements and non-stationarity [20, 21]. Finally, recent work has extended GPs to RNA-seq data using a negative binomial likelihood [22]. Together, these advances establish a rich foundation for the methods introduced in this paper.

This work makes three primary advances in the application of Gaussian processes to sparse, irregularly sampled longitudinal metabolomics data. First, we develop and rigorously compare via simulations two variable selection strategies: a structured step-wise search algorithm and a continuous shrinkage approach employing a Horseshoe prior on kernel hyperparameters to identify relevant biomarkers with high sensitivity and specificity. Second, we apply one of our methods to real-world data as an example, identifying known as well as plausible novel biomarkers. Third, we integrate these modeling innovations into `waveome`, an open-source Python library that incorporates hyperparameter tuning, enables inducing point approximations for computational scalability, and generates visualizations and diagnostic metrics.

## 2 Results

### 2.1 Waveome achieves superior recovery of complex latent dynamics

To evaluate the performance of our proposed methods, we simulated observations from four distinct data-generating processes, chosen to reflect both the diversity and realism of longitudinal biological measurements. These include random intercepts (capturing subject-specific baselines), periodicity (cyclical or seasonal variation), linear trends (gradual directional changes), and higher-order interactions (nonlinear and rapidly fluctuating dynamics). Collectively, these processes span a range of functional forms; some are relatively simple and can be well-approximated by standard approaches such as linear regression or mixed-effects modeling, while others require more flexible, nonparametric representations. Detailed specifications for each process are provided in Section 4.5. Figure 2 illustrates representative draws under two random seeds, showing latent functions (top rows) and corresponding negative binomial count observations (bottom rows), which highlight both the structured patterns and stochastic variability present in the data.

**Fig. 2:**
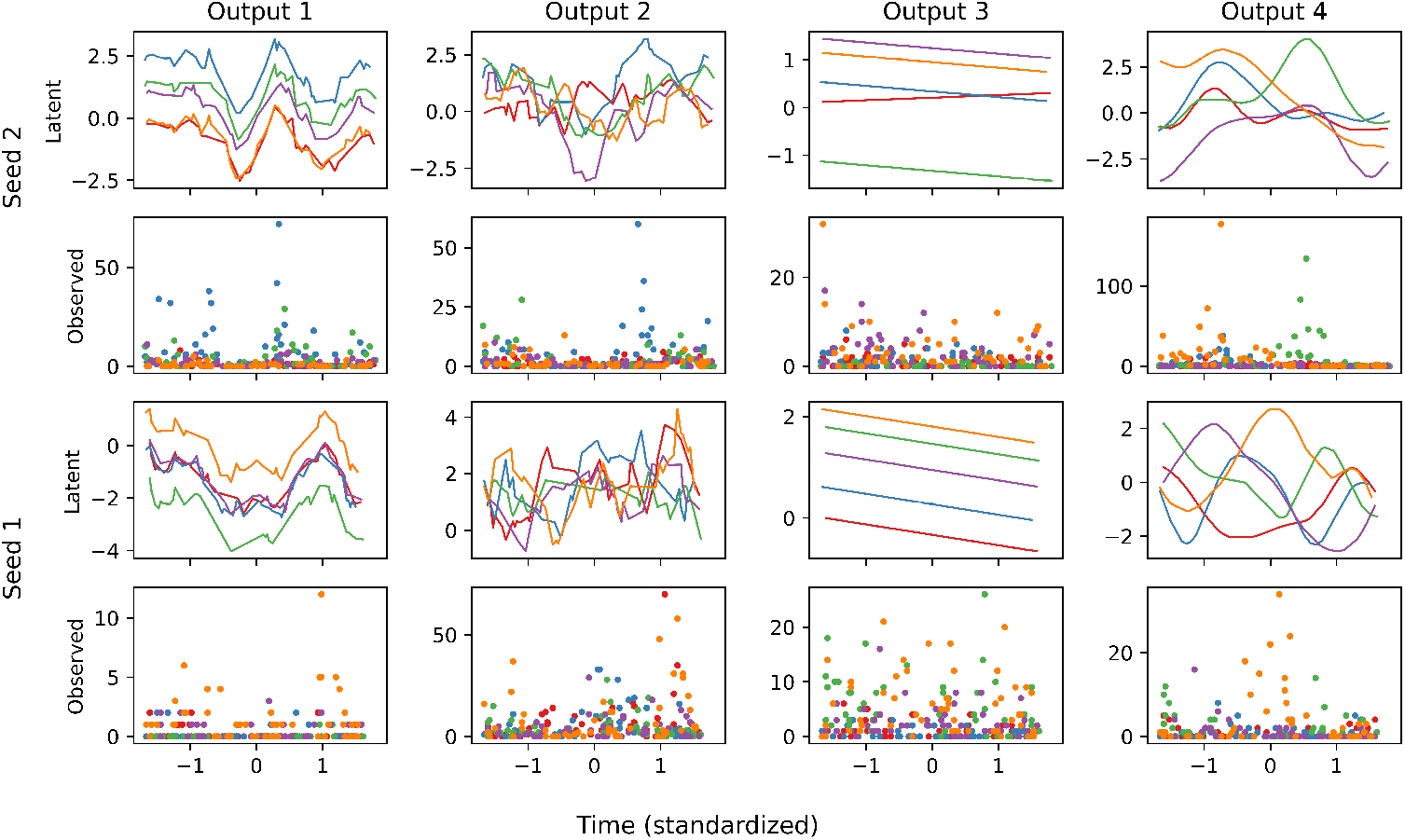
Example draws from our simulation framework illustrating four distinct latent processes and their corresponding count observations under a negative binomial likelihood. Each column corresponds to one of the four kernel-defined outputs, while in each block, the top subpanel displays five latent function realizations and the bottom subpanel shows the (sampled) generated counts. The upper two rows use one seed while the lower two rows use another, demonstrating the stochasticity present with fixed simulation parameters. Parameters used for the figures were *N* = 5, *n*_*i*_ = 50, *ϵ* = 0, and *α* = 1.

We evaluated both `waveome` strategies against six benchmarks: two linear models on log-transformed data (mixed-effects regression and LASSO), two generalized models with a negative binomial likelihood (GLM and GAM), and two Gaussian process baselines employing Automatic Relevance Determination (ARD) with and without a negative binomial likelihood (NB-ARD) [23]. Across 50 simulation replicates per parameter set, GP-based methods with a negative binomial likelihood—namely `waveome` (both search and penalization) and NB-ARD—achieved the lowest holdout KL-divergence (Figure 3), demonstrating superior distributional fit. Notably, NB-ARD exhibited greater variability in performance compared to the more stable `waveome` approaches.

**Fig. 3:**
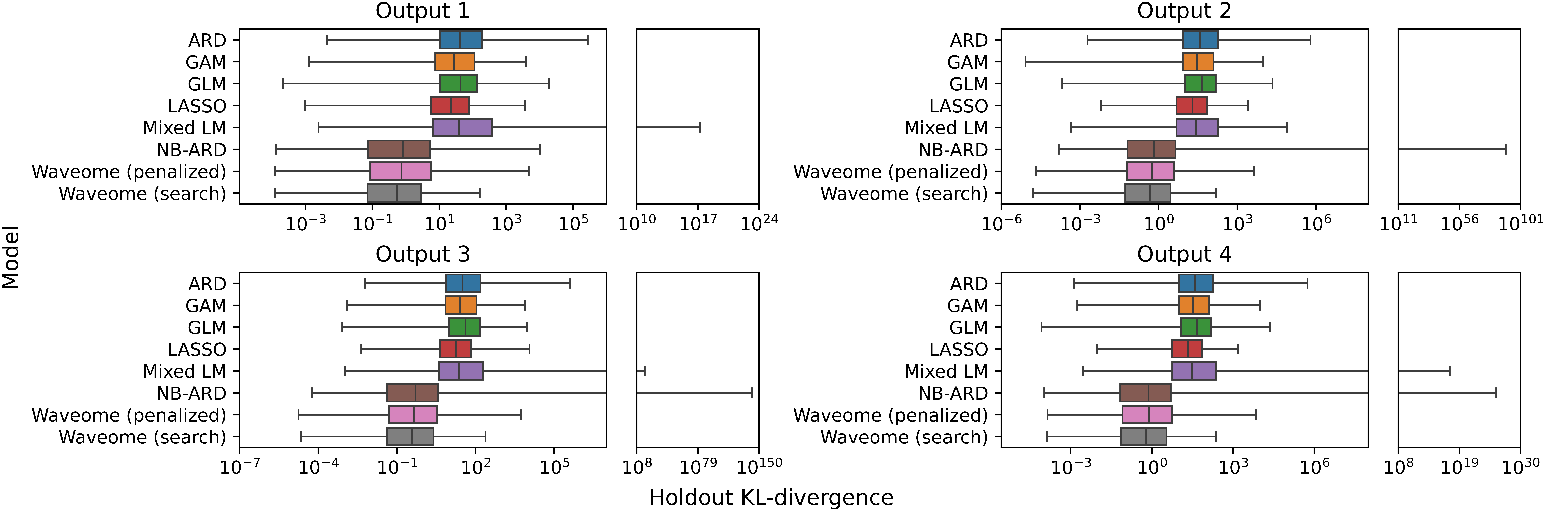
Holdout KL-divergence values for each model type across 50 simulation replicates, evaluated on four simulated outcomes. Lower values indicate better agreement between the model’s predictive distribution and the true data-generating process. Box-plots display the inter-quartile range while the whiskers depict the full range, with extreme outliers separated in the right-hand subpanels for clarity. Gaussian process models with a negative binomial likelihood, including `waveome` penalized and search variants, consistently achieve lower divergence compared to linear and standard GP baselines. The NB-ARD model, while competitive in median performance, displays substantially higher variability relative to `waveome` methods.

Feature selection performance is summarized in Figure 4. An ideal method with no Type I or Type II errors would appear in the top-right corner, combining high sensitivity with high specificity. In this setting, specificity corresponds to 1 − *α*, where *α* is the Type I error rate (false positives), and sensitivity corresponds to 1 − *β*, where *β* is the Type II error rate (false negatives). The penalization-based `waveome` method offered the best balance, maintaining sensitivity comparable to ARD variants while achieving higher specificity, thereby reducing false positives. The search-based variant was the most conservative, attaining the highest specificity but at the cost of lower sensitivity, highlighting the trade-off between detecting all relevant features and avoiding spurious selections.

**Fig. 4:**
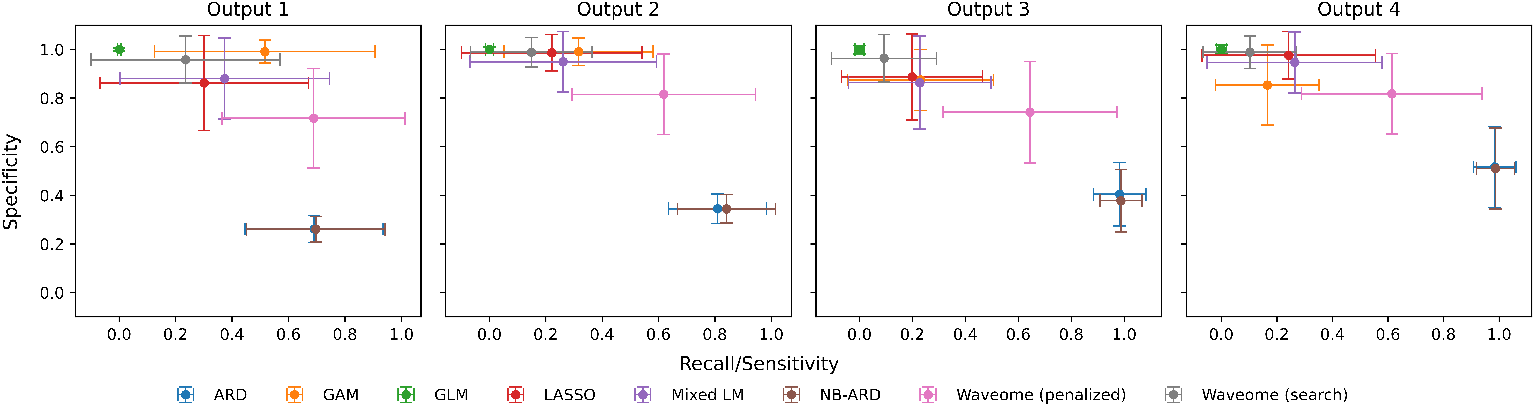
Recall (sensitivity) and specificity for feature selection across eight modeling approaches, averaged over 50 simulation replicates for four simulated outcomes. Points show mean values, and error bars denote one standard deviation. Specificity corresponds to 1 − *α* (avoiding Type I errors) and sensitivity to 1 − *β* (avoiding Type II errors). The penalized `waveome` approach combines strong sensitivity with higher specificity than ARD-based methods, reducing false positives. The search-based variant and linear models maximize specificity but with reduced sensitivity.

### 2.2 Metabolic signatures track clinical severity in Crohn’s disease

We applied the penalized `waveome` framework to Crohn’s disease samples from the NIH iHMP IBD cohort [24], retaining 238 observations per metabolite across 564 labeled metabolites. The penalized approach is favored in this context, given the higher sensitivity and our interest in identifying potential biomarkers for further study. Example trajectories for mandelate (Figure 5) highlight irregular sampling and skewed distributions. Seventy-two metabolites showed significant associations with Harvey–Bradshaw Index (HBI) scores using both linear and squared exponential kernels. We show the top twenty metabolites based on log Bayes factor in Figure 6. These metabolites clustered into six functional classes: short-chain fatty acids, bile acids, acylcarnitines, amino acids/proteolytic metabolites, lipids, and other small molecules. Posterior functions for lithocholate are shown in Figure 7.

**Fig. 5:**
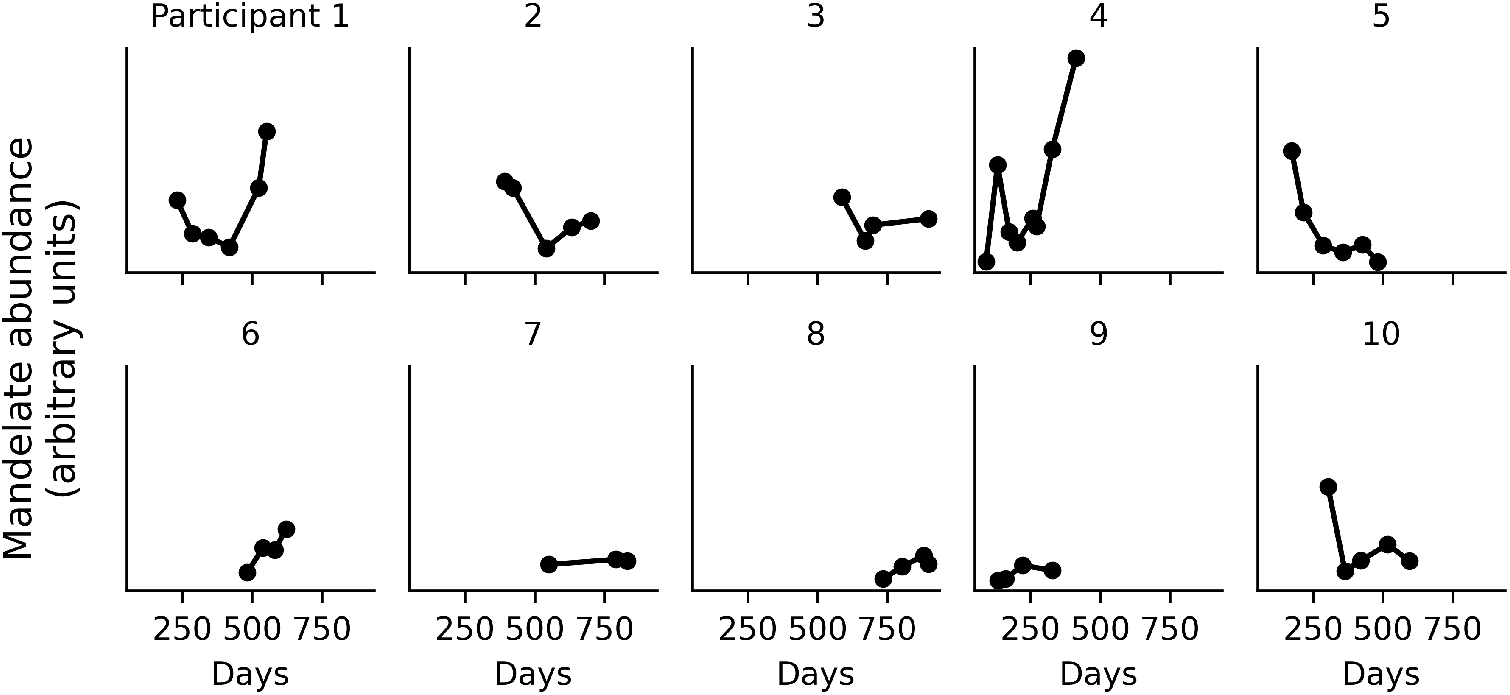
Longitudinal mandelate intensity trajectories for ten Crohn’s disease patients in the NIH Integrative Human Microbiome Project. Each panel shows measured mandelate levels over study days for a single individual; both the number and timing of observations differ across participants. Intensities are highly right-skewed, illustrating challenges in modeling sparse, irregularly sampled metabolomic time series.

**Fig. 6:**
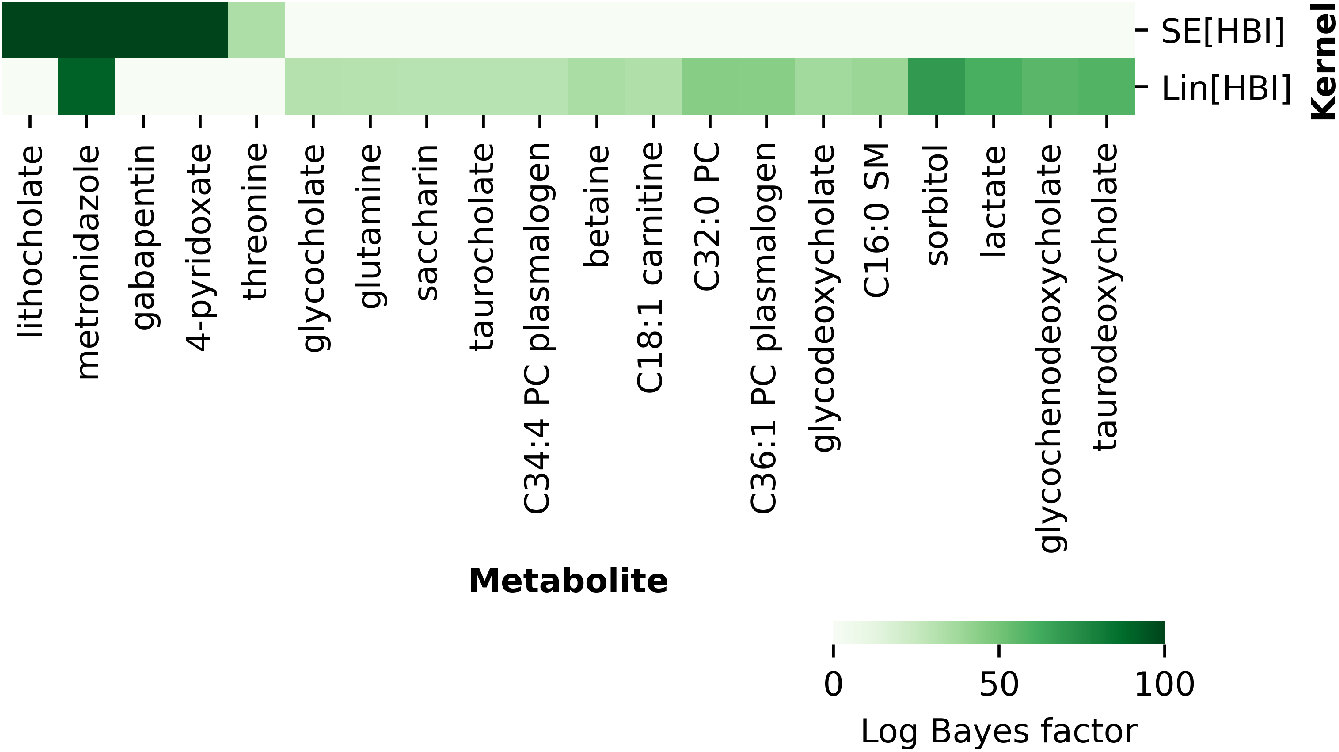
Heatmap of top twenty metabolite cross-sectional models based on log Bayes factor, with selected linear (Lin) and squared exponential (SE) kernels governed by HBI levels. Color indicated log Bayes factor for given kernel component in final metabolite model.

**Fig. 7:**
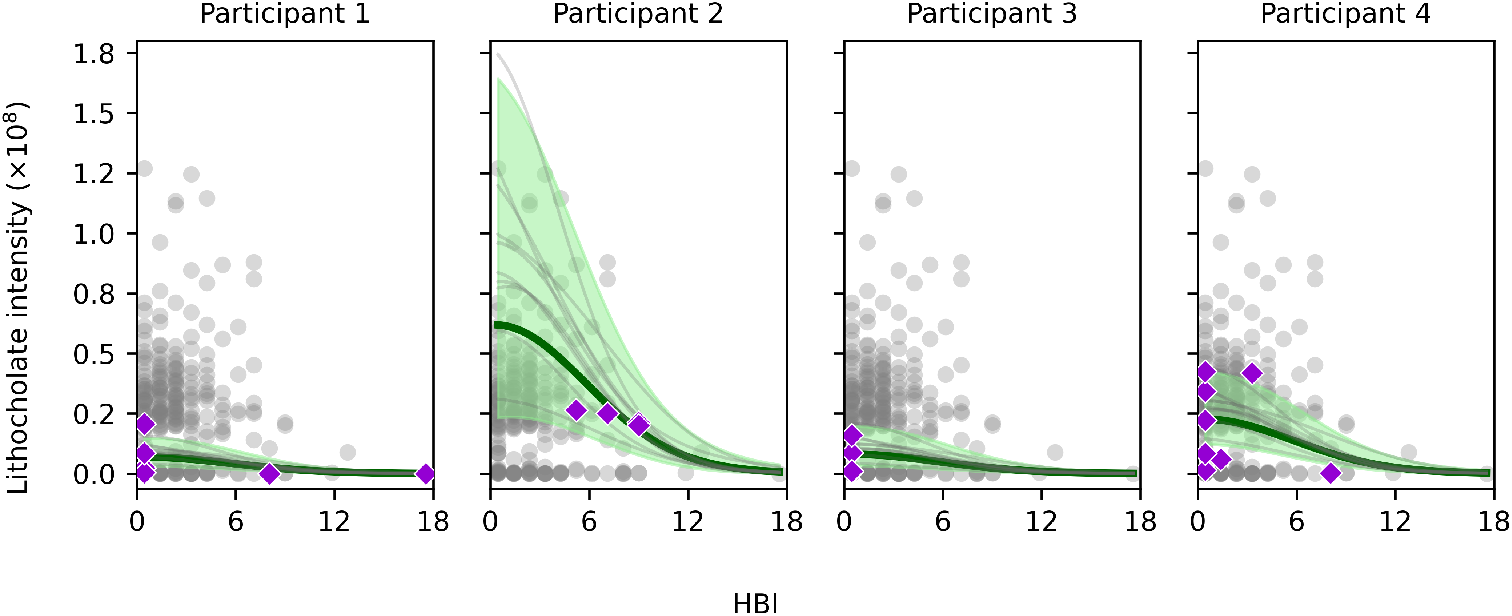
Posterior function mean and sample draws from the fitted lithocholate model, conditional on four participants in the cohort represented in each facet. Participant observations are indicated by the purple diamonds compared to the entire sample in the scatterplot, dark green signifies the average lithocholate intensity, light green bands are the 95% confidence intervals, and gray lines are sample draws. The resulting model shows an individual-specific intercept offset as well as a common squared exponential HBI kernel where higher values of HBI are associated with lower lithocholate intensity values.

Using metadata-aligned trajectories, we modeled metabolite dynamics around each participant’s peak HBI value. Nine metabolites exhibited significant temporal associations (Table 1), spanning bile acids (e.g., taurolithocholate), lipid species (specific TAGs, cholesteryl esters), and other small molecules (oxalate, betaine, 4-methylcatechol). An example decomposition for oxalate is shown in Figure 8.

**Table 1:**
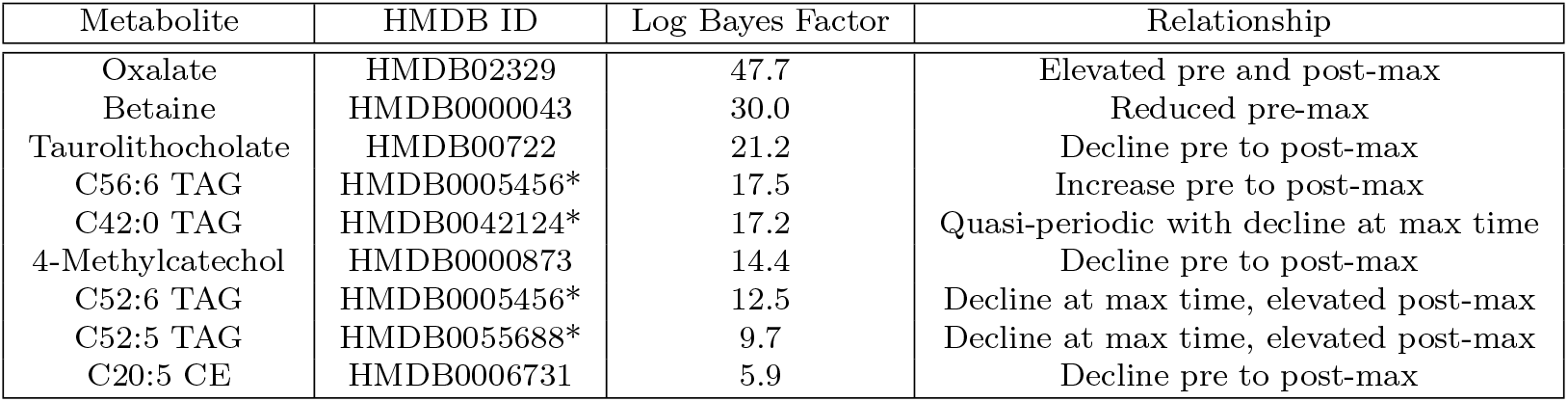
Metabolites selected as relevant for temporal associations with max severity time with the corresponding log Bayes factor and a characterization of the dynamic profile around the max sample point.

| Metabolite | HMDB ID | Log Bayes Factor | Relationship |
| --- | --- | --- | --- |
| Oxalate | HMDB02329 | 47.7 | Elevated pre and post-max |
| Betaine | HMDB0000043 | 30.0 | Reduced pre-max |
| Taurolithocholate | HMDB00722 | 21.2 | Decline pre to post-max |
| C56:6 TAG | HMDB0005456* | 17.5 | Increase pre to post-max |
| C42:0 TAG | HMDB0042124* | 17.2 | Quasi-periodic with decline at max time |
| 4-Methylcatechol | HMDB0000873 | 14.4 | Decline pre to post-max |
| C52:6 TAG | HMDB0005456* | 12.5 | Decline at max time, elevated post-max |
| C52:5 TAG | HMDB0055688* | 9.7 | Decline at max time, elevated post-max |
| C20:5 CE | HMDB0006731 | 5.9 | Decline pre to post-max |

**Fig. 8:**
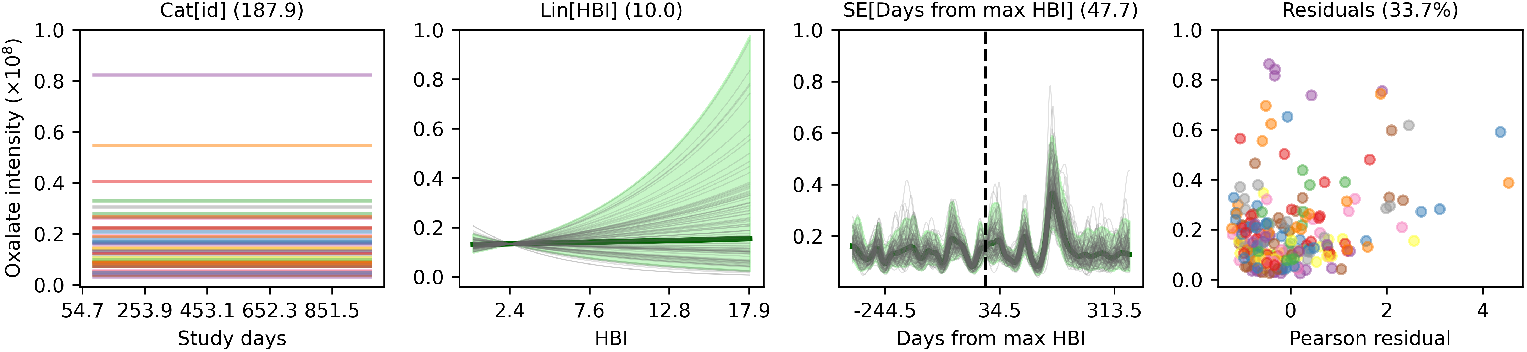
Marginal posterior predictions from the fitted oxalate model. Each facet is a kernel component with a corresponding log Bayes factor in the parenthesis, except the last, which shows the residuals of the model with the overall model’s unexplained deviance. The first component shows an individual average offset, second shows a linearly increasing relationship between HBI and oxalate levels, and the third visualizes the dynamics around the max HBI point showing elevated levels around the event. The dashed vertical line is the point of max severity per individual. The final facet shows the Pearson residuals by observed oxalate values, colored by participant ID.

## 3 Discussion

This paper presents two complementary Gaussian process–based methods for analyzing sparse, irregularly sampled longitudinal omics data with automatic variable selection: a structured kernel search and a penalization approach using a Horseshoe prior. Through simulation benchmarking, we demonstrated that both `waveome` variants consistently outperform linear, generalized additive, and standard GP baselines in distributional accuracy as measured by KL-divergence, with differing trade-offs in sensitivity and specificity. The penalization method prioritized sensitivity in variable selection, whereas the search-based strategy emphasized specificity, illustrating that the two approaches can be tuned to match differing research priorities.

Applying `waveome` to the NIH iHMP Crohn’s disease cohort uncovered both cross-sectional and temporal metabolite associations. Cross-sectionally, we identified six functional metabolite classes—short-chain fatty acids (SCFAs), bile acids, acylcarnitines, amino acids/proteolytic metabolites, lipids, and other compounds—that were significantly associated with disease severity. Several of these classes are well supported by the literature: SCFAs and bile acids are consistently depleted or altered in Crohn’s disease [25, 26], amino acid–derived metabolites (e.g., tryptophan catabolites) and acylcarnitines have been repeatedly reported as enriched in CD stool [25, 27], and sphingolipid and complex lipid perturbations are increasingly recognized as features of IBD-associated dysbiosis [25, 28]. In contrast, many detected metabolites in the “other compounds” category—such as certain vitamins, simple sugars, and xenobiotics—lack clear mechanistic links to CD pathophysiology, highlighting areas for further investigation.

The biological plausibility of these associations is supported by known mechanisms: reduced SCFAs compromise gut barrier integrity and promote inflammatory signaling [29–31], altered bile acid pools exacerbate mucosal inflammation through signaling disruption [32], elevated acylcarnitines may foster dysbiosis and augment immune responses [27, 33], and amino acid deficiencies impair mucosal healing and immune function [34]. Disturbances in lipid metabolism, including phosphatidylcholines and sphingomyelins, can alter cell membrane composition and downstream signaling cascades [35]. Moreover, certain medications (e.g., metronidazole) and dietary components (e.g., sorbitol) identified in our models can influence the microbiome and inflammation, potentially confounding or mediating observed associations.

In addition to cross-sectional findings, `waveome` detected temporal associations aligned to peak disease activity, identifying nine metabolites with significant dynamic changes. These include the bile acid taurolithocholate and several lipid species (C56:6 TAG, C42:0 TAG, C52:5/6 TAGs, C20:5 CE) as well as oxalate, betaine, and 4-methylcatechol. Oxalate and bile acid dysregulation are well-established features of CD pathophysiology [26, 36–42]. The TAG and cholesteryl ester species we report have been observed in untargeted lipidomics of IBD [43, 44] but lack mechanistic validation; they may reflect broad fat malabsorption or shifts in microbial lipase activity. Betaine has been linked to methylation and osmoprotective roles in gut epithelial cells [45], and 4-methylcatechol is a microbial polyphenol catabolite with putative anti-inflammatory effects [25, 46]. Neither has been longitudinally characterized in relation to CD flares, making them promising targets for follow-up studies.

While `waveome` captures non-linear temporal dependencies and performs robust variable selection, the current implementation assumes omic feature independence, and it models metabolite intensities without explicit mechanistic constraints. Future extensions could: (i) incorporate zero-inflated or hurdle likelihoods to better distinguish between undetectable and absent metabolites; (ii) refine kernel search and penalization hyperparameter tuning via heuristic or Bayesian optimization to reduce computational cost; (iii) introduce multivariate GP priors to model metabolite co-regulation and enable pathway-level inference; and (iv) embed causal inference frameworks (e.g., instrumental variables, time-varying interventions) to distinguish correlation from causation, clarifying whether observed associations reflect upstream drivers or downstream consequences (such as medication use). By addressing these limitations, `waveome` can evolve into an even more powerful platform for longitudinal biomarker discovery, mechanistic hypothesis generation, and integrative multi-omics modeling. We encourage the biomedical research community to adopt, critique, and contribute to its ongoing development.

## 4 Methods

### 4.1 Gaussian Processes

A GP defines a distribution over functions *f* : *χ* → ℝ such that for any finite collec-tion of inputs 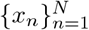, the vector of function values **f** = [*f* (*x*_1_), …, *f* (*x*_*N*_)]^T^ follows a multivariate normal distribution [12]. A GP is fully specified by its mean function *m*(*x*) and covariance (kernel) function *k*(*x, x*^*′*^), both governed by a vector of hyperparameters *θ*. Hence, a GP denoted by *f* (*·*) ~ 𝒢 𝒫_*θ*_ (*m*(·), *k*(·, ·)) where

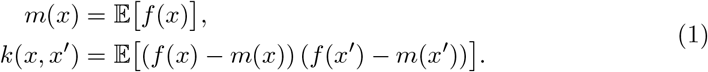

In practice, one often sets *m*(*x*) ≡ 0 and models observations 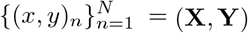 with additive Gaussian noise:

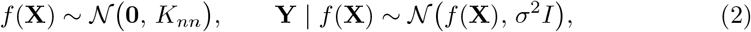

where (*K*_*nn*_)_*ij*_ = *k*_*θ*_(*x*_*i*_, *x*_*j*_). The selection of the kernel function is critical, as it encodes prior assumptions about smoothness, periodicity, or other functional properties [47], and thus constrains the space of admissible functions (see Section 4.2 for definitions and Figure 9 for illustrative draws under different kernels).

**Fig. 9:**
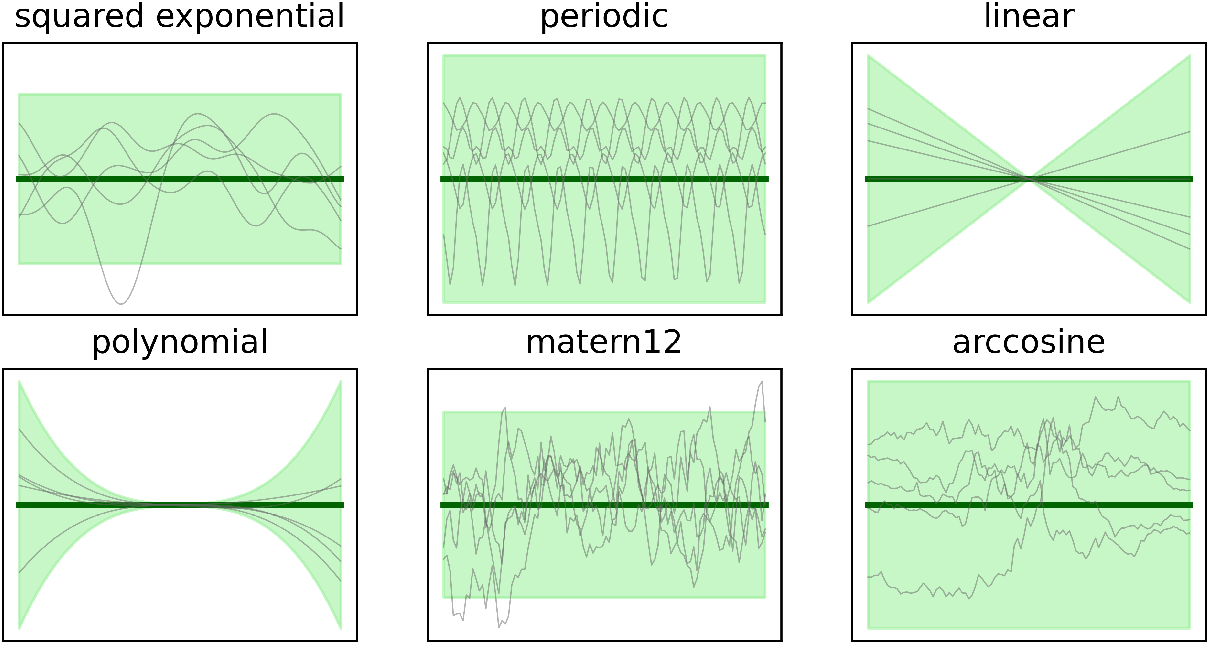
Prior function draws under six distinct Gaussian process kernels, illustrating how kernel choice shapes the space of supported functions. Each panel corresponds to a different covariance function, displaying multiple sample paths (gray) and the 95% prior confidence interval (green). The dark green line denotes the prior mean (here set to zero), highlighting differences in smoothness, periodicity, growth behavior, and roughness induced by each kernel.

The hyperparameters *θ* of the kernel and the noise variance *σ*^2^ are typically chosen by maximizing the log marginal likelihood of the data

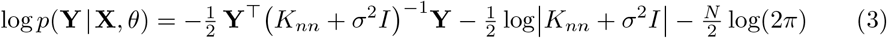

either using first or second-order optimization methods, depending on the size of the parameter space. The log marginal likelihood decomposes into three parts on the right-hand side of (3): the fit to the observed data, model complexity, and a normalizing constant. The first term measures how well the observations conform to the current covariance assumption, or in other words a kernel-weighted measure of residual error. The second term penalizes complexity, rising when the GP prior becomes diffuse over many possible datasets, and so discourages overly flexible covariance choices. The third term is a constant normalization that depends only on the sample size.

Maximizing the marginal likelihood therefore balances these forces: the data-fit term drives the model to explain the observations (reducing bias), while the complexity penalty prevents excessive flexibility (controlling variance). In practice, changing hyperparameters alters both the data-fit and complexity penalty, producing the familiar bias–variance trade-off. Figure 10 contrasts posterior predictions under default versus optimized hyperparameters for a squared-exponential kernel.

**Fig. 10:**
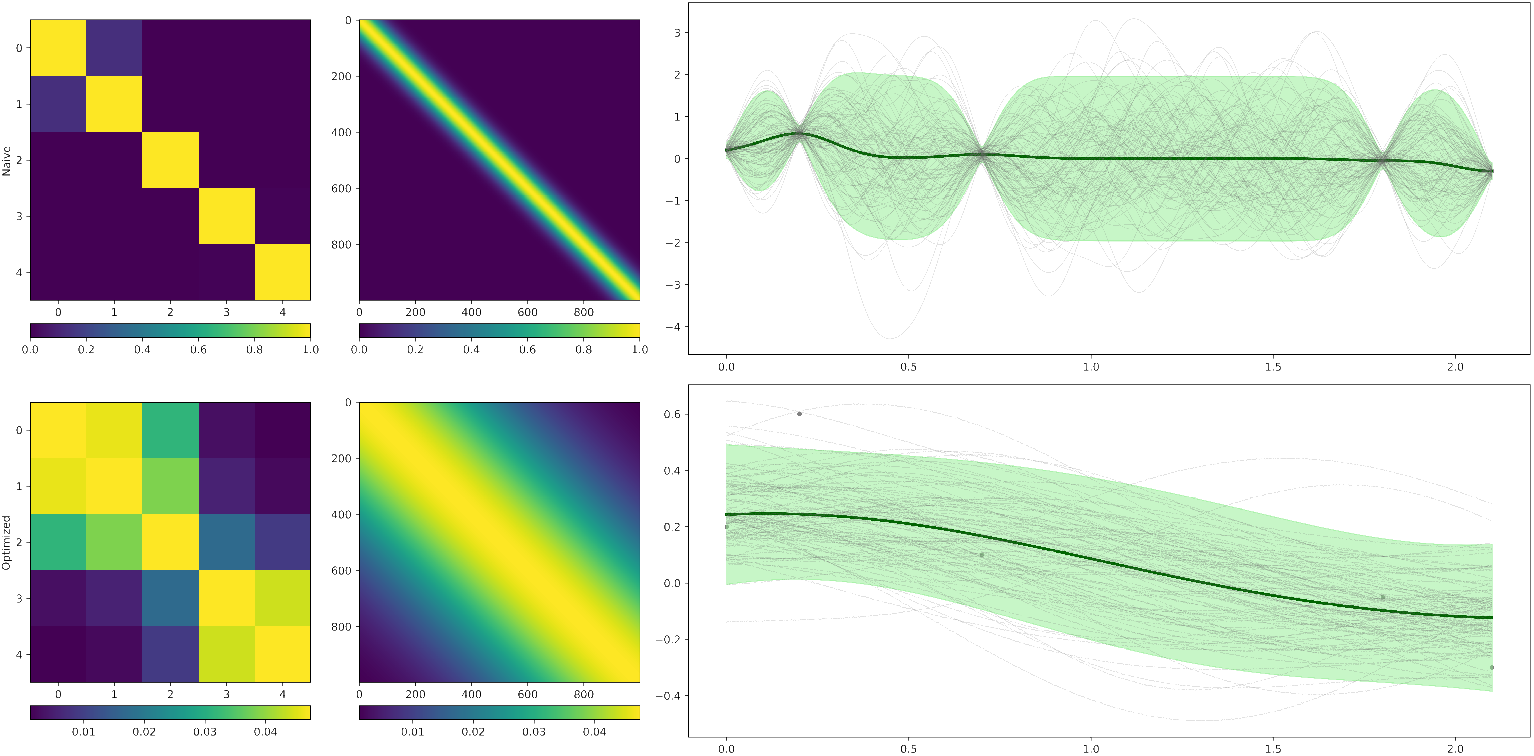
Impact of hyperparameter optimization on Gaussian process predictions with a squared-exponential kernel. The top row illustrates the unoptimized model: the left panel shows the prior covariance matrix among five observed inputs, the center panel displays the prior covariance over 1,000 equally spaced test points, and the right panel depicts the resulting posterior samples (gray), posterior mean (dark green), and 95% confidence intervals (light green). With default hyperparameters, the model exhibits weak correlations and high uncertainty away from observations. The bottom row shows the same panels after maximizing the marginal likelihood: the optimized covariance matrices reflect increased correlation length and variance, and the posterior mean closely tracks the data with substantially reduced uncertainty, demonstrating how hyperparameter tuning critically shapes GP inference.

A powerful feature of GPs is closure under addition and multiplication – sums and products of valid kernels produce new valid kernels. This property enables structured priors, such as additive decompositions or interaction terms, that can reflect hierarchical effects. To accommodate repeated measurements or other categorical covariates, we specify a categorical kernel which yields a compound-symmetric covariance structure and acts analogously to a random intercept when added to other kernels. Moreover, the product of a categorical kernel with a time-based squared-exponential kernel produces individual-specific smooth trajectories

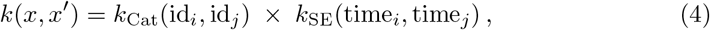

thereby capturing both between-subject variability and within-subject temporal dynamics.

### 4.2 Kernel functions

The covariance (kernel) functions in our GP models can be seen in Table 2. Here, *σ*^2^ is the variance parameter, 𝓁 is the lengthscale, *γ* is the period, *τ* is the offset, *d* is the polynomial degree, and 𝟙{·} is the indicator function.

**Table 2:**
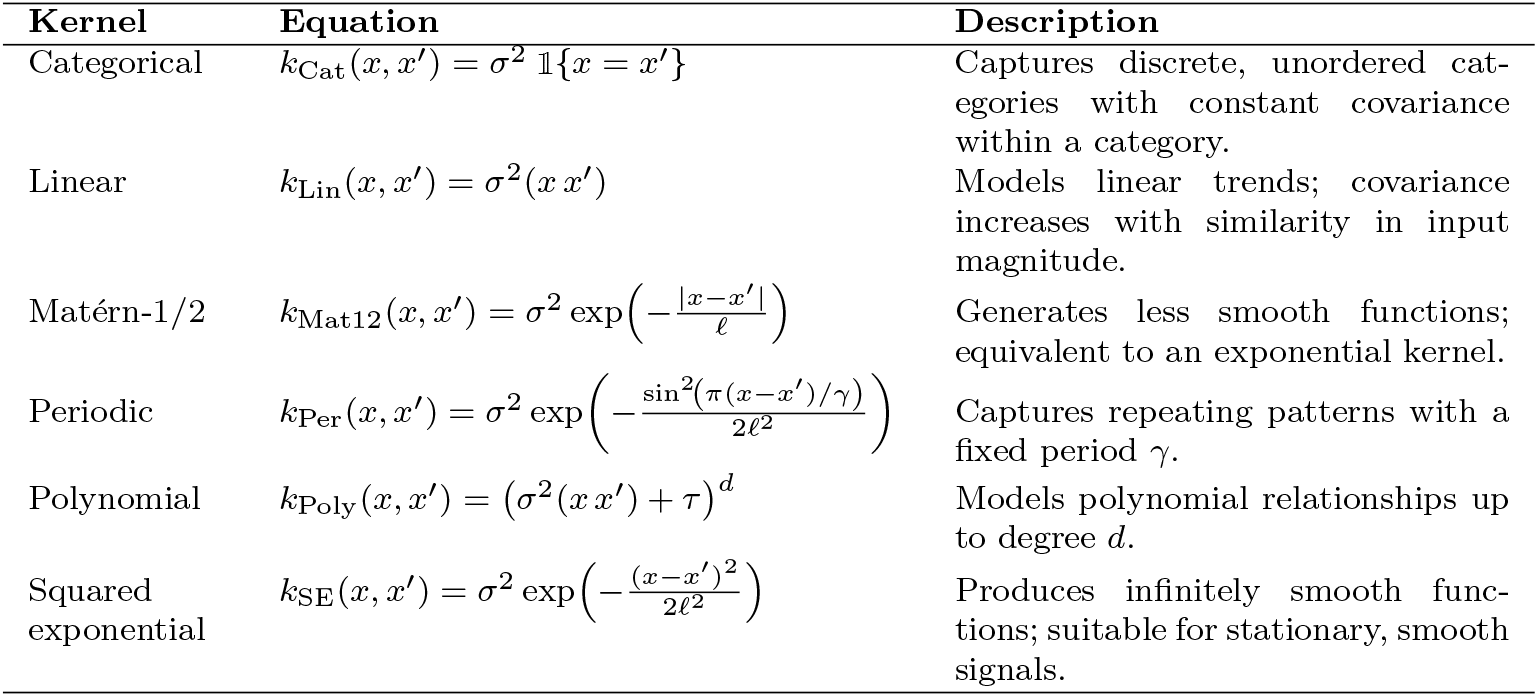
Covariance (kernel) functions used in Gaussian process models, with corresponding mathematical definitions and brief descriptions.

### 4.3 Model Extensions

We incorporate three key extensions to the standard GP framework to achieve both computational scalability and flexibility for longitudinal count data. First, we employ an inducing point approximation that projects the full covariance matrix *K*_*nn*_ onto a lower-dimensional subspace spanned by *m* ≪*n* inducing inputs. We replace *K*_*nn*_ with the low-rank surrogate

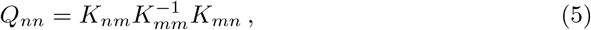

reducing the Cholesky decomposition cost from 𝒪 (*n*^3^) to 𝒪 (*n*^2^*m*) [48–50]. The inducing locations may be fixed to a subset of observed inputs or treated as variational parameters and optimized jointly with the kernel hyperparameters.

Second, to accommodate non-Gaussian likelihoods, specifically the negative binomial model for over-dispersed metabolite intensities, we adopt a variational inference scheme. Because the marginal log likelihood is now intractable, we assume a Gaussian variational posterior

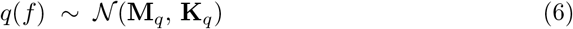

over the latent function values *f*, and we optimize the evidence lower bound (ELBO)

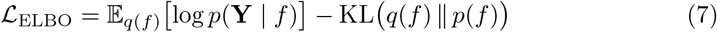

with respect to both the variational parameters and the kernel hyperparameters [22, 51–53]. This framework seamlessly integrates with the inducing-point approximation to maintain scalability under complex likelihoods.

Finally, we leverage stochastic optimization via mini-batching and adaptive gradient methods within the `TensorFlow` ecosystem [54–56]. By computing unbiased estimates of the ELBO gradient on small random subsets of the data, we further reduce per-iteration cost and memory requirements, making it feasible to fit high-dimensional longitudinal models from large datasets. All of these components, inducing points, variational non-Gaussian inference, and scalable stochastic optimization, are implemented in `GPflow` [57, 58], and exposed through our `waveome` library for streamlined longitudinal biomarker discovery.

### 4.4 Variable Selection

Nonparametric Gaussian process models offer exceptional flexibility but can become computationally burdensome and overparameterized when faced with many covariates.

This is especially true when there are multiple kernels to consider per variable and hierarchical structures in the data, such as repeated sampling. To address this, we introduce two strategies for variable selection within the GP framework: a kernel-space search and a penalization approach.

#### Search-based Kernel Selection

Our search method performs a forward stepwise exploration of kernel combinations, using the Bayesian Information Criterion (BIC) to guide selection. We begin by fitting separate GPs for each continuous covariate *x*_*i*_ paired with every candidate kernel 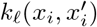, and for each categorical covariate using the categorical kernel *k*_cat_. The model with the lowest BIC using a single kernel term initializes the search. At each subsequent iteration, we expand the current best model(s) by adding or multiplying one new kernel component (e.g., 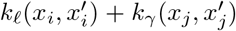) and optimize the new GP’s hyperparameters. New models whose BIC falls within a predefined tolerance (e.g.ΔBIC ≤ 6) of the best score are retained for the next round. We continue this process until no improvement in BIC is observed or a maximum search depth is reached (see Figure 11 for an illustration). This procedure efficiently identifies parsimonious kernel structures that balance goodness-of-fit and model complexity.

**Fig. 11:**
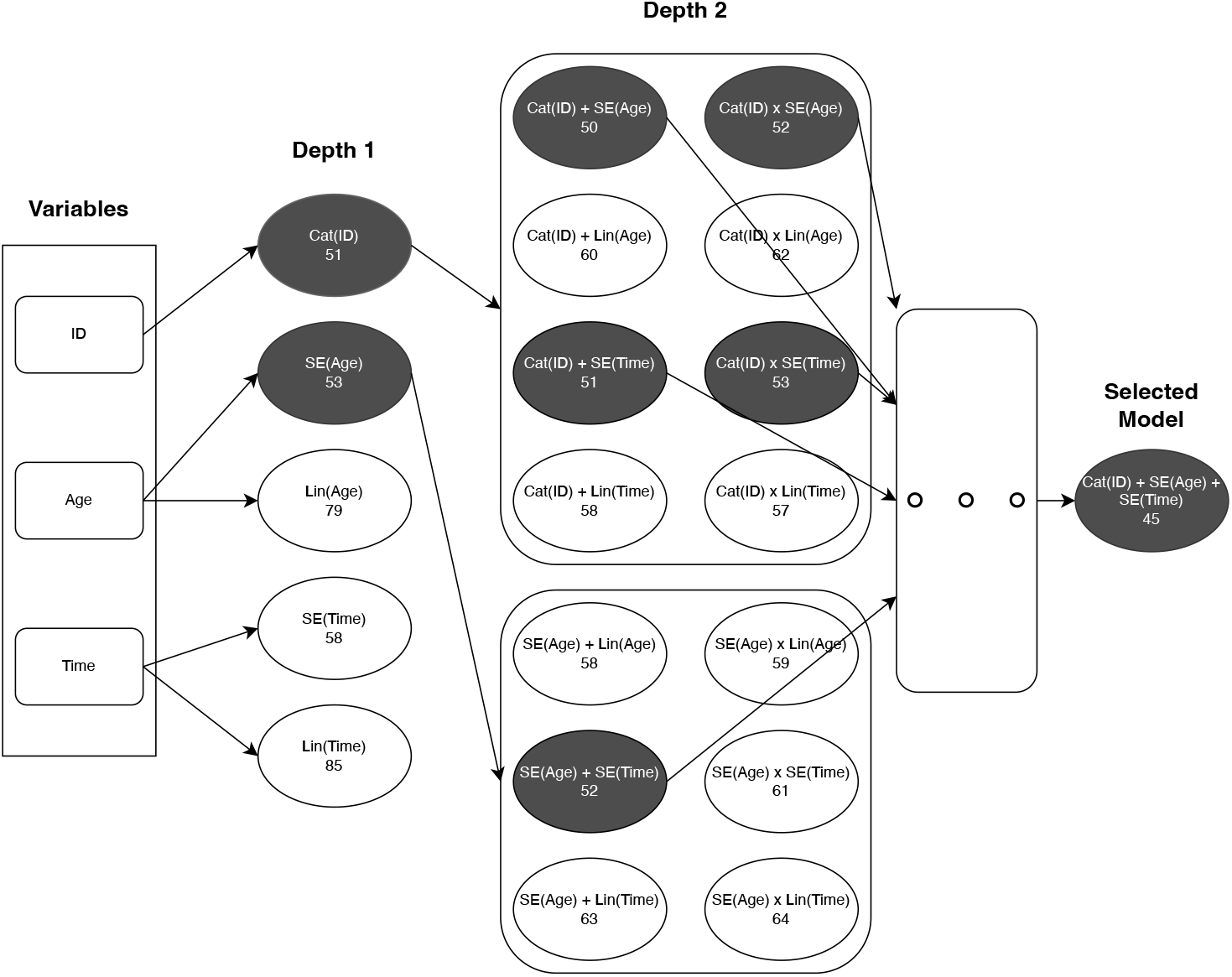
An example with three variables using forward stepwise kernel-space search for Gaussian process variable selection using BIC. At depth 1, univariate models are fit for each covariate, unit ID with a categorical (Cat) kernel, and age and time each paired with squared exponential (SE) or linear (Lin) kernels. The number instead each node is its BIC score. At depth 2, new models are generated by adding (+) or interacting (×) one additional kernel component to the best depth 1 configurations. Nodes whose BIC lies within ΔBIC ≤ 6 of the optimal score are carried forward (highlighted), and this process repeats until no further improvement is made. The final selected model appears at the far right.

#### Penalization

As an alternative to discrete kernel-space search, we construct a “fully saturated” GP prior that includes all main-effect kernels and up to second-order interactions, then impose a shrinkage penalty on each kernel’s variance hyperparameter. Whereas L1 regularization (the Laplace prior) applies a uniform shrinkage rate across parameters, the Horseshoe prior adapts to the signal strength of each component through a two-level hierarchy

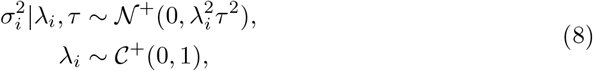

where 𝒩^+^ is a half-Gaussian distribution and 𝒞^+^ is a half-Cauchy distribution. Here, the global parameter *τ* encourages overall sparsity, while each local parameter *λ*_*i*_ allows an individual kernel component to avoid heavy shrinkage if it carries a strong signal. In contrast, L1 regularization enforces the same decay rate on all components, which can be problematic when combining stationary and nonstationary kernels in the same model. However, the Horseshoe automatically adapts to each 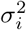’s magnitude, ensuring that spurious effects are effectively driven to zero, yet high-variance kernels retain the flexibility needed to capture broad trends. This yields a parsimonious, mixed-scale GP model without manual tuning of individual kernel penalty strengths.

### 4.5 Simulation study

#### Setup

To assess model fit and variable selection performance in controlled settings, we generated longitudinal count data from four known GP priors under a range of sampling and noise regimes. These included varying numbers of units *N* ∈ {10, 50, 100, 500}, rate per unit *λ* ∈ {2, 4, 8, 16}, and latent noise 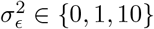 injected. The simulated latent GP values were then passed through a negative binomial likelihood with varying amounts of dispersion *α* ∈ {1, 5, 10} as another simulation parameter. The complete data-generating process for unit *i*, time *j*, and output *k* is

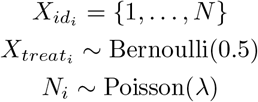

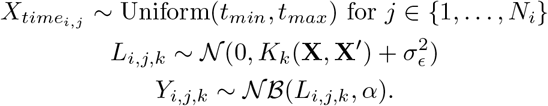

Each of the four latent GP models is governed by kernels *K*_*k*_ that are tied to at least two of the three input variables: individual ID 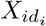, a binary treatment flag 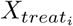, and a time point 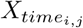, which are then concatenated to produce the full input matrix **X**. The four kernels, with their hyperparameters, are as follows:

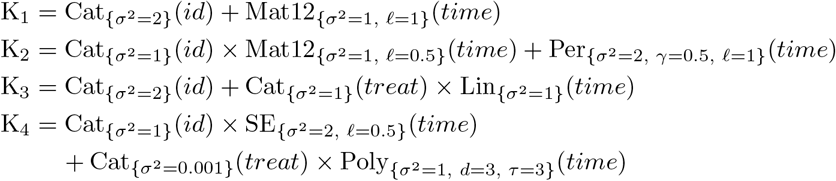

#### Comparable methods

To benchmark `waveome` we compared its two variants (search and penalization) to six commonly used alternatives: a linear mixed-effects model (Mixed LM), LASSO regression (LASSO), a negative-binomial generalized linear model (GLM), a negative-binomial generalized additive model (GAM), Gaussian process regression with ARD on a Gaussian likelihood (ARD), and a variational GP with ARD and a negative-binomial likelihood (NB-ARD). For all comparators we followed a consistent high-level pipeline: (i) apply method-appropriate preprocessing, (ii) perform variable or model selection according to each method’s standard practice, and (iii) obtain holdout predictions for evaluation. We summarize the inputs and selection procedures in Table 3, while more detailed information can be found in the Supplementary information below.

**Table 3:** High-level summary of comparator methods (inputs and variable-selection rules). Detailed implementation is provided in the Supplementary Methods.

| Method | Inputs | Variable selection |
| --- | --- | --- |
| Mixed LM | $\log(y+1)$; subject ID and time random effects | Nested random/fixed-effect comparison |
| LASSO | $\log(y + 1)$; one-hot IDs, interactions, standardized covariates | Cross-validated LASSO penalty and nonzero coefficients |
| GLM | Raw intensities; one-hot IDs, interactions | dispersion tuning; retain significant terms |
| GAM | Raw intensities; one-hot IDs, spline basis for time | Smoothness-term tuning; retain significant terms |
| ARD | $\log(y + 1)$; one-hot IDs, interactions, standardized covariates, SE kernels | ARD lengthscales (relevance via lengthscale) |
| NB-ARD | Raw intensities; one-hot IDs, interactions, standardized covariates, SE kernels | ELBO optimization; ARD lengthscales (relevance via lengthscale) |

### 4.6 iHMP study cohort

We analyzed stool metabolomics from the Inflammatory Bowel Disease Multi-omics Database (IBDMDB / iHMP) cohort [24], accessed via the project homepage (https://ibdmdb.org/). The iHMP study collected longitudinal multi-omics measurements and clinical metadata (including Harvey–Bradshaw Index, HBI) on participants with inflammatory bowel disease across multiple study sites; here we restricted analyses to participants with a clinical diagnosis of Crohn’s disease (CD). HBI was used as the primary disease-severity measure in cross-sectional and temporal analyses [59].

#### Data selection and preprocessing

From the iHMP metabolomics tables we analyzed only labeled metabolite features. We imposed a missingness filter and kept features for which at least 50% of available samples had a measured value greater than zero; after this filter, 564 labeled metabolites remained for analysis. Across the retained features, there were 238 observations per metabolite in total (median 5 observations per individual). All preprocessing steps were kept deliberately minimal to preserve observed intensity structure and to reflect realistic, over-dispersed peak-area measurements. Metabolite measurements were treated as continuous outcomes in downstream modeling and undetected values were imputed as zero. The candidate covariates considered in models were: HBI score, days from the individual’s maximum HBI, participant identifier (ID), study site, race, sex, time in study (absolute study day), age, and participant self-reported general well-being.

#### Significance criterion and reproducibility

For the `waveome` penalized approach applied to the iHMP data, a metabolite was deemed to show a significant association with a covariate if the corresponding kernel variance parameter remained above 10^−4^ after optimization. Each metabolite model was initialized with a fully saturated kernel. Specifically, for continuous covariates (HBI score, days from maximum HBI, time in study, age) we included both squared exponential and linear kernels, and for categorical covariates (participant ID, study site, race, sex, general well-being) we included a categorical kernel. Model hyperparameters were randomly initialized and optimized using the L-BFGS-B algorithm until convergence. All random seeds, exact initialization settings, optimization options, and the scripts used to construct, fit, and evaluate the penalized models are provided in the project repository.

## Acknowledgments

The authors would like to thank A.R. Taheriyoun for testing the software and reviewing the manuscript. We also thank the GW Research Computing team for access to the Pegasus high-performance computing cluster (Pegasus: https://hpc.gwu.edu/pegasus/). This work was supported by the National Science Foundation award number 2109688 to A.R.

## Author contributions

A.R. conceived and supervised the study and secured the funding that supported this work. A. Ross developed the software, contributed to experimental design and implementation, prepared the data, and conducted the applied analyses. J.L.-P. contributed to methodological development and manuscript editing. All authors interpreted the results, contributed to drafting the manuscript, and approved the final version.

## Code availability

The `waveome` source code, including all Gaussian process modeling routines, variable selection implementations, simulation scripts, and example workflows used in this study, is openly available at https://github.com/omicsEye/waveome. The repository contains installation instructions, documentation, and reproducible scripts for replicating the analyses presented in this manuscript. `waveome` is released under the MIT License.

## Supplementary information

### Detailed implementation of comparable methods

All code used for the benchmarks and the analyses described below is available in the project repository. The text below documents computational resources, exact preprocessing, selection rules, parameter grids, and fallback strategies used in the simulation benchmarks.

### Computational resources

Simulation experiments were performed on the George Washington University high performance computing cluster Pegasus. For the simulation runs we used Pegasus’ medium-memory compute nodes: Dell PowerEdge R740 servers (dual 20-core Intel Xeon Gold 6148 processors, up to 40 physical cores per node), 384 GB DDR4 RAM, 800 GB local SSD, and Mellanox EDR InfiniBand interconnect (100 Gb/s). Jobs were submitted as batch jobs and typically requested 8 CPU cores and the required memory per task.

### Common preprocessing

An 80/20 train/holdout split was applied to each simulated dataset. For methods assuming Gaussian errors (Mixed LM, LASSO, ARD) the response was transformed as log(*y* + 1). Predictions from these methods were evaluated using a Gaussian likelihood for KL-divergence metrics. Subject identifiers and other categorical covariates were encoded either as categorical factors (for `statsmodels` formula interfaces) or one-hot encoded (drop-first) when using matrix-based solvers (e.g., `scikit-learn` pipelines). Continuous covariates were standardized (zero mean, unit variance) where required by the method. If solver issues occured, the training mean and standard deviation were used for holdout predictions as a fallback.

### Mixed LM (linear mixed-effects models)

- Software: `statsmodels.mixedlm`.
- Candidate random effect structures: `1 + time + treat`, `1 + time`, `1 + treat`, `1`, and fixed-effects only.
- Selection rule: nested model comparisons via likelihood improvements, choosing the
- most parsimonious model that yields a statistically significant improvement.
- Predictive distribution: Gaussian with mean as point estimate prediction and standard deviation from model residuals.

### LASSO (L1-penalized linear model)

- Software: `scikit-learn` (`LassoCV`, `PolynomialFeatures`, `StandardScaler`).
- Preprocessing: response log(*y* + 1), one-hot encoding for categorical IDs (drop-first), interaction-only polynomial features for covariates, standardization of design matrix.
- Selection rule: cross-validated LASSO penalty tuning; features with absolute coefficient > 0.01 considered selected.
- Predictive distribution: Gaussian with mean as point estimate prediction and standard deviation from model residuals.

### GLM (negative binomial GLM)

- Software: `statsmodels.formula.api.glm` with `NegativeBinomial` family.
- Dispersion tuning: grid search over dispersion parameter *α* ∈ {1, …, 10} with selection by maximum log-likelihood.
- Variable selection: initial saturated model (subject, time, treatment, interactions)
- followed by Wald tests; terms with *p* < 0.05 retained for the reduced model.
- Robustness: use alternative optimizers or clustered covariances when standard fits fail; fallback to training mean otherwise.
- Predictive distribution: negative binomial mean from point estimate prediction and
- dispersion parameter from the final fitted model.

### GAM (negative binomial GAM)

- Software: `statsmodels.gam.GLMGam` with `BSplines` basis.
- Spline specification: B-spline on time with degrees of freedom and degree chosen (e.g., `df=12`, `degree=3`).
- Dispersion tuning: grid search *α* ∈ {1, …, 10} as for `glm`.
- Term selection: Smoothness penalty selected based on cross-validation; Wald tests on parametric terms (*p* < 0.05).
- Prediction caveat: holdout times clipped to the training time range to avoid knot extrapolation issues.
- Predictive distribution: negative binomial mean from point estimate prediction and dispersion parameter from the final fitted model.

### ARD (GP with Gaussian likelihood and ARD)

- Software: `GPflow` with squared-exponential kernel supporting ARD (one lengthscale per input).
- Preprocessing: same encoding/standardization pipeline as for LASSO; response transformed as log(*y* + 1).
- Relevance inference: lengthscale magnitudes interpreted as relevance; empirical threshold used in scripts (lengthscale within (0.01, 6) considered “relevant”).
- Predictive distribution: Gaussian with a posterior predictive mean and the estimated standard deviation from residuals parameter.

### NB-ARD (Variational GP with NB likelihood and ARD)

- Software: `GPflow` with a variational GP implementation and the project’s negative binomial likelihood wrapper.
- Preprocessing: same design matrix as LASSO/ARD; raw counts modeled directly.
- Inference: variational Gaussian posterior with ELBO optimization; ARD length-scales used for relevance assessments with the same empirical cutoff strategy as ARD.
- Predictive distribution: negative binomial with a posterior predictive mean and an estimated dispersion parameter.

